# Development of a Porcine Cell Line Stably Expressing Ephrin-B2 for Nipah Virus Research and Diagnostic Testing

**DOI:** 10.1101/2025.09.30.679668

**Authors:** Hui Zhang, Akatsuki Saito

**Author notes:** China Animal Health and Epidemiology Center, Qingdao, Shandong 266032, China (Hui Zhang).

## Abstract

Nipah virus (NiV) is a highly pathogenic zoonotic virus transmitted from bats to humans through pigs as a crucial intermediate host. NiV outbreaks pose significant public health and economic threats, especially for pig farmers. Although the World Organization for Animal Health (WOAH) recommends African green monkey-derived Vero cells for NiV isolation, pig-derived cell lines could represent an optimal platform for propagating pig-origin NiV, as signs and symptoms of NiV infection differ among different hosts. In this study, we generated and evaluated pig-derived PK-15 cells stably expressing pig-derived ephrin-B2 (PK-15/Ephrin-B2 cells), the primary receptor for NiV. NiV pseudovirus infectivity was increased by >1000-fold in PK-15/Ephrin-B2 cells compared with that in wild-type PK-15 cells, whereas virus susceptibility was higher in PK-15/Ephrin-B2 cells than in Vero cells (>30-fold). Furthermore, *Stat2*-knockout PK-15/Ephrin-B2 cells exhibited stable viral infectivity in the presence of type I interferon, making it particularly suitable for clinical sample testing. Moreover, PK-15/Ephrin-B2 cells proved useful for neutralization tests using anti-NiV hyperimmune ascitic fluid. Therefore, PK-15 cells expressing pig ephrin-B2 could represent an efficient tool for virus isolation, vaccine development, and virological studies of NiV and related henipaviruses.

## 1. Introduction

Nipah virus (NiV), a single negative-stranded RNA virus belonging to the genus *Henipavirus* of the family *Paramyxoviridae*, can infect pigs [1], leading to significant economic losses in the swine industry [2]. Pigs can also serve as intermediate hosts for NiV between bats and humans. In the Malaysian outbreak, more than one million pigs were culled, and economic losses reached as high as 400 million US dollars [2]. To date, 265 people have been infected through direct contact with NiV-infected pigs, resulting in 105 deaths [3, 4]. In other NiV outbreaks, although pigs lacked clinical signs, 44.2% of pigs in the outbreak area were anti-NiV antibody-positive [5]. These findings indicate that pigs play an essential role in the spread of NiV infection, highlighting the importance of controlling NiV infection in pigs.

Ephrin-B2 and ephrin-B3 are the only identified receptors for NiV [6–8]. These receptors are highly conserved across Mammalia, thereby explaining the broad species tropism of NiV [9]. During outbreaks, NiV isolates from pigs share high sequence similarity with those from humans [10]. However, the manifestation of infection differs between humans and pigs. In the Malaysian outbreak, NiV caused severe-to-fatal central nervous system infections in humans, suggesting that the central nervous system is the most severely affected by NiV [11]. Conversely, NiV infection in pigs leads to severe respiratory and neurological symptoms. Lesions appear mainly in the lungs, brain, and kidneys. Furthermore, a large proportion of pigs exhibit asymptomatic infection [12]. Therefore, the viral tropism of NiV and clinical outcome of NiV infection can differ depending on the infecting host species.

The endothelial and smooth muscle cells of small blood vessels are the primary targets of NiV and the regions responsible for histologic changes in humans [13]. Therefore, various cell lines, mostly originating from humans, have been evaluated for NiV permissibility [14–16]. Given the differences in the respiratory symptoms in pigs, porcine airway epithelial primary cells have also been used to evaluate NiV infection [17–19]. Although the World Organization for Animal Health (WOAH) recommends African green monkey-derived Vero cells for NiV isolation [20], pig-derived cell lines could represent an optimal platform for propagating pig-origin NiV, as the signs and symptoms of infection differ among infected hosts despite the virus’s broad host range.

Furthermore, during viral isolation, contamination by type I interferon (IFNs) or IFN-inducible substances such as double-stranded RNA or lipopolysaccharides can exert antiviral effects in the target cells. In this cascade, both IFN alpha and beta receptor subunit 1 (IFNAR1) on the cell surface and intracellular STAT2 play critical roles [21]. We recently generated the pig-derived PK-15 cell line lacking *Ifnar1* and *Stat2* to overcome this limitation [22].

In this study, we established PK-15 cells stably expressing pig Ephrin-B2 (PK-15/Ephrin-B2 cells) to improve their susceptibility of NiV that utilizes ephrin-B2. A viral infection assay using an HIV-1–based viral vector pseudotyped with NiV G and F proteins illustrated that the NiV pseudovirus displayed >1000-fold higher infectivity in PK-15/Ephrin-B2 cells than in normal PK-15 cells. Furthermore, we demonstrated the usefulness of PK-15 cells lacking *Stat2* with stable expression of pig ephrin-B2. Finally, we demonstrated that anti-NiV hyperimmune ascitic fluid specifically neutralized NiV pseudovirus in PK-15/Ephrin-B2 cells. These findings suggest that PK-15/Ephrin-B2 cells are promising tools for isolating and propagating NiV and other ephrin-B2–dependent viruses from pig samples.

## 2. Materials and methods

### 2.1 Plasmids

The psPAX2-IN/HiBiT and pWPI-Luc2 plasmids were kindly gifted by Dr. Kenzo Tokunaga [17]. pMD2.G was a gift from Dr. Didier Trono (Cat# 12259; http://n2t.net/addgene:12259, accessed on May 10, 2025; RRID: Addgene_12259).

### 2.2 Cell culture

Lenti-X 293T (TaKaRa, Kusatsu, Japan, Cat# Z2180N), PK-15 (Japanese Collection of Research Bioresources Cell Bank, Ibaraki, Japan, Cat# JCRB9040), and PK-15 *Stat2* knockout (*Stat2* k/o) cells [22] were cultured in Dulbecco’s modified Eagle’s medium (Nacalai Tesque, Kyoto, Japan, Cat# 08458-16) supplemented with 10% fetal bovine serum and 1× penicillin–streptomycin (Nacalai Tesque, Cat# 09367-34).

### 2.3 Generation of a Retroviral Vector to Express Ephrin-B2

To generate a retroviral vector expressing pig ephrin-B2, the coding sequence of pig ephrin-B2 was synthesized using the amino acid sequences deposited in GenBank (accession number: ABV44489.1) with codon optimization to pig cells (Integrated DNA Technologies, Inc., Coralville, IA, USA). The synthesized DNA sequence is summarized in File S1. Synthesized DNA was cloned into the pDON-5 Neo-vector (TaKaRa, Kusatsu, Japan, Cat# 3657), which was prelinearized by *Not*I-HF (New England Biolabs, Ipswich, MA, USA, Cat# R3189L] and *Bam*HI-HF (New England Biolabs, Cat# R3136L) using an In-Fusion HD Cloning Kit (TaKaRa, Cat# Z9633N). Plasmids were amplified using 5-alpha F′ *Iq* competent *Escherichia coli* (New England Biolabs, Cat# C2992H) and extracted using the PureYield Plasmid Miniprep System (Promega, Madison, WI, USA, Cat# A1222). The plasmid sequence was verified using a SupreDye v3.1 Cycle Sequencing Kit (M&S TechnoSystems, Osaka, Japan, Cat# 063001) with a Spectrum Compact CE System (Promega).

### 2.4 Generation of PK-15 Cells Stably Expressing Ephrin-B2

Lenti-X 293T cells were co-transfected with pDON-5 Neo-pig ephrin-B2 plasmid, pGP packaging plasmid (TaKaRa, Cat# 6160), and pMD2.G plasmid with TransIT-293 Transfection Reagent (TaKaRa, Cat# V2700) in Opti-MEM (Thermo Fisher Scientific, Waltham, MA, USA, Cat# 31985062). The supernatant was filtered 2 days after transfection. Collected retroviral vectors were used to infect normal PK-15 or PK-15 *Stat2* k/o cells, which were then cultured in 500 µg/mL G-418 (Nacalai Tesque, Cat# 09380-44) for 6 days. Single-cell cloning was then performed with an SH800S cell sorter (Sony, Minato-Ku, Japan). Briefly, a single cell clone was sorted into one well of a 96-well plate. After cell growth, ephrin-B2 expression in each clone was evaluated by western blotting.

### 2.5 Preparation of plasmids for expressing NiV G and F proteins

The codon-optimized NiV G (accession number: AAK29088.1) and F genes (accession number: AAK29087.1) were separately inserted into the pTwist vector and synthesized by Twist Bioscience (South San Francisco, CA, USA). G protein had a C-terminal HA tag (YPYDVPDYA), and F protein had a C-terminal AU1 tag (DTYRYI). None of the tags affected G and F protein function, as previously described [23].

### 2.6 Production of HIV-1-based and MLV-based viral vectors pseudotyped with NiV G and F proteins

To rescue an HIV-1–based viral vector pseudotyped with NiV G and F proteins (NiV pseudovirus), Lenti-X 293T cells (2.5 × 10^5^ cells/mL) were co-transfected with 400 ng of psPAX2-IN/HiBiT, 400 ng pWPI-Luc2, 40 ng of pTwist-NiV-G, and 160 ng of pTwist-NiV-F plasmids using 3 μL of TransIT-293 Transfection Reagent in 100 μL of Opti-MEM I Reduced Serum Medium. To rescue the MLV-based pseudovirus, the same procedure was conducted, albeit with co-transfection with 400 ng of pGP, 400 ng of pDON-5 Neo-luc2, 40 ng of pTwist-NiV-G, and 160 ng of pTwist-NiV-F plasmids. The supernatant was filtered 2 days after transfection. The cell lysate was collected to evaluate G and F protein expression by western blotting.

### 2.7 Virus Infection

Wild-type PK-15 cells and PK-15/Ephrin-B2 #3 cells were seeded into 96-well plates at 1 × 10^4^ cells/well, as previously described [22]. To test viral infectivity in the presence of type I IFN, cells were treated with 25 units/mL Universal Type I IFN (PBL Assay Science, Piscataway, NJ, USA, Cat# 11200) for 24 h. For the luciferase-encoding virus, 10 μL of virus solution were added to each well. Infected cells were lysed 2 days after infection using britelite plus (PerkinElmer, Shelton, CT, USA, Cat# 6066769), and the luminescent signal was measured using a GloMax Explorer Multimode Microplate Reader (Promega).

### 2.8 Neutralization Test

PK-15/Ephrin-B2 #3 cells were seeded into a 96-well plate at 1 × 10^4^ cells/well and incubated for 24 h. Pseudotyped NiV solution was mixed with different dilutions of polyclonal anti-NiV hyperimmune mouse ascitic fluid (BEI Resources, Cat# NR-48961). After incubation at 37℃ for 1 h, 20 μL of the mixture was added to each well, and the luminescent signal was measured 2 days after infection using a GloMax Explorer Multimode Microplate Reader.

### 2.9 Western Blotting

To evaluate ephrin-B2 expression, pelleted cells were lysed in 2× Bolt LDS sample buffer (Thermo Fisher Scientific, Cat# B0008) containing 2% β-mercaptoethanol (Bio-Rad, Hercules, CA, USA, Cat# 1610710) and incubated at 70°C for 10 min. The expression of NiV G, NiV F, and ephrin-B2 was evaluated using SimpleWestern Abby (ProteinSimple, San Jose, CA, USA) with anti-HA tag (6E2) mouse monoclonal antibody (Cell Signaling, Danvers, MA, USA, Cat# 2367S), anti-AU1 epitope tag antibody (BioLegend, San Diego, CA, USA, Cat# 901905), and anti-Myc tag rabbit monoclonal antibody (Cell Signaling Biotechnology, Cat# 2272S, ×125), along with the Anti-Mouse Detection Module (ProteinSimple, Cat# DM-001) and Anti-Rabbit Detection Module (ProteinSimple, Cat# DM-001). The amount of input protein was visualized using the Total Protein Detection Module (ProteinSimple, Cat# DM-TP01). The expected sizes of HA-tagged NiV G, AU1-tagged NiV F, and Myc-tagged ephrin-B2 are 68.1, 62.1, and 38.1 kDa, respectively, according to the Protein Molecular Weight website (https://www.bioinformatics.org/sms/prot_mw.html, accessed on May 10, 2025).

### 2.10 Alignment of Ephrin-B2 Protein Sequences

The MUSCLE algorithm on MEGA-X was used to align the ephrin-B2 protein sequences from pigs (Acc. Num: ABV44489.1) and humans (Acc. Num: AAH69342.1). The alignment was visualized using CLC Genomics Workbench software (version 22.0.1, QIAGEN, Hilden, Germany).

### 2.11 Phylogenetic Analysis of Mammalian Ephrin-B2

A phylogenetic tree was constructed using ephrin-B2 sequences obtained from GenBank for various mammals including pigs (accession number: ABV44489.1), humans (accession number: AAH69342.1), cattle (accession number: XP_019827252.2), horses (accession number: ABV44488.1), dogs (accession number: ABV44487.1), cats (accession number: ABV44486.1), mice (accession number: AAH57009.1), chimpanzees (accession number: JAA26050.1), gorillas (accession number: XP_004054766.1), orangutans (accession number: XP_054302618.1), bears (accession number: XP_026349366.1), bats (accession number: XP_054430356.1), rabbits (accession number: XP_069904683.1), elephants (accession number: XP_049723308.1), whales (accession number: XP_068382904.1), sheep (accession number: XP_004012283.1), and camels (accession number: XP_010984339.1). The tree was created using the neighbor-joining method with the Jones–Taylor–Thornton model implemented in a MEGA X program.

### 2.12 Calculation of the Identity of Ephrin-B2 among Animal Species

The identity of ephrin-B2 sequences among animal species was calculated using MEGA X with a pairwise distance matrix. Analyses were conducted using the Poisson correction model. The rate variation among sites was modeled with a gamma distribution (shape parameter = 5). All ambiguous positions were removed for each sequence pair (pairwise deletion option).

### 2.13 Statistical Analysis

The results were presented as the mean and standard deviation of six measurements from one assay, representing at least two or three independent experiments. Differences in infectivity between two different conditions (e.g., between PK-15 and PK-15/Ephrin-B2 #3 cells) were evaluated using an unpaired, two-tailed Student’s *t*-test. *p* ≤ 0.05 was considered significant. Tests were performed using Prism 9 software v9.1.1 (GraphPad, Boston, MA, USA).

## 3. Results

### 3.1 Generation of PK-15 Cells Stably Expressing Pig Ephrin-B2

We constructed a phylogenetic tree using mammalian ephrin-B2 amino acid sequences retrieved from GenBank, thereby revealing genetic differences in ephrin-B2 sequences between humans and pigs (**Figure 1a**). The homology rate of ephrin-B2 sequences among different animals was approximately 94%, with the rate increasing to approximately 97% between humans and pigs (**Figure 1b**), suggesting that NiV has the potential to infect a wide range of mammals. However, as signs and symptoms differ among animals, it was necessary to establish a pig-derived cell line that can highly support viral infection and replication.

**Figure 1.**
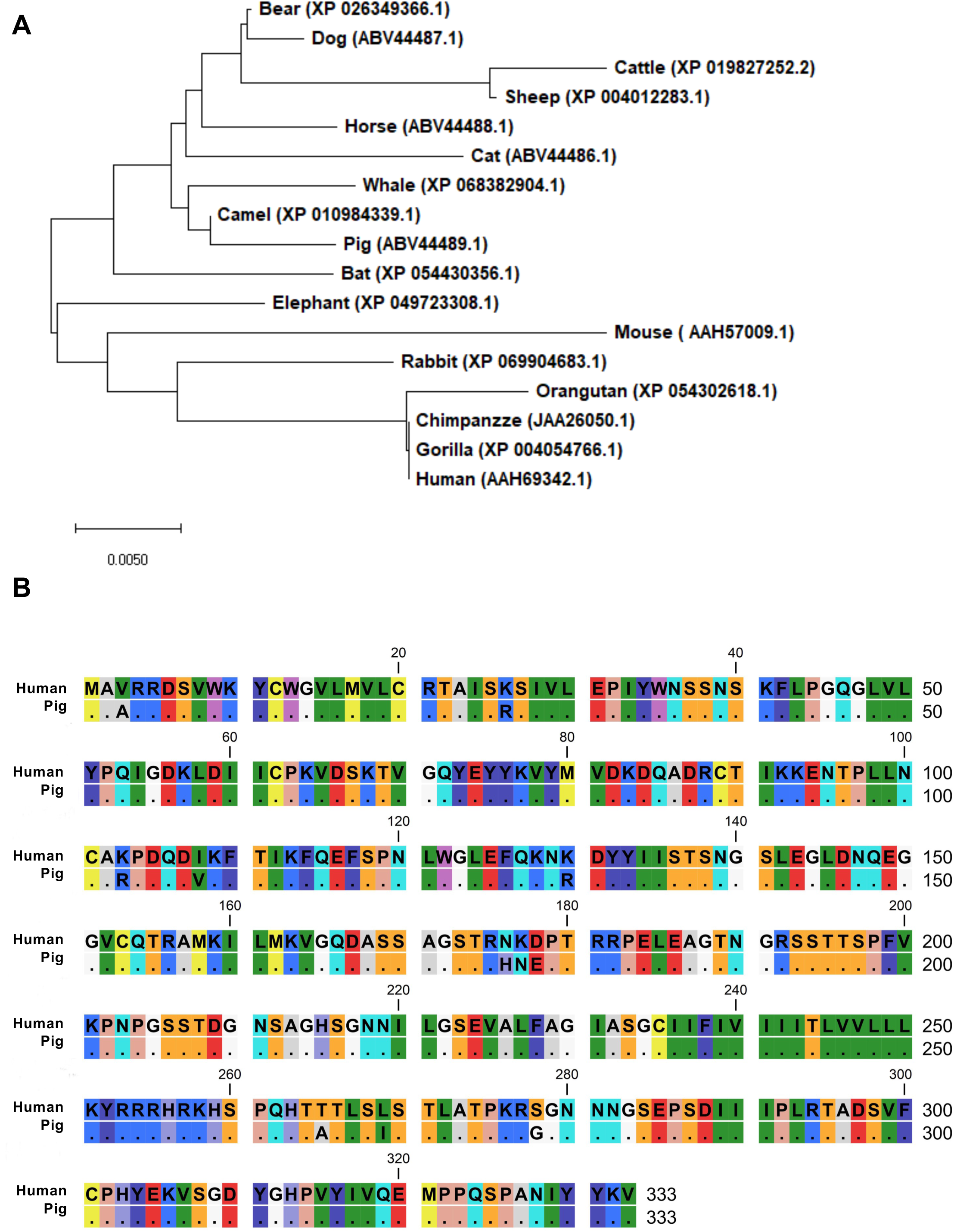
Genetic diversity of mammalian ephrin-B2. (a) A phylogenetic tree was constructed using ephrin-B2 sequences retrieved from GenBank. The tree was constructed using the neighbor-joining method with the Jones–Taylor–Thornton model implemented in MEGA X. (b) Amino acid alignment of ephrin-B2 sequences between humans and pigs.

Therefore, we designed Myc-tagged pig ephrin-B2 using pig genomic information deposited in GenBank (accession number: ABV44489.1) with pig codon optimization. The synthesized DNA was cloned into a retroviral vector, which was used to infect normal PK-15 cells, and the transduced cells were selected with neomycin. Single-cell clones were obtained via cell sorting, and western blotting using anti-Myc tag antibody was performed on each cell clone to test pig ephrin-B2 expression (**Figure 2a**). We selected PK-15/Ephrin-B2 clone #3 because it exhibited higher Ephrin-B2 expression than the other clones. Furthermore, we expressed pig ephrin-B2 in PK-15 cells lacking *Stat2* [22], thereby obtaining PK-15 (*Stat2* k/o)/Ephrin-B2 #19 cells (**Figure 2b**).

**Figure 2.**
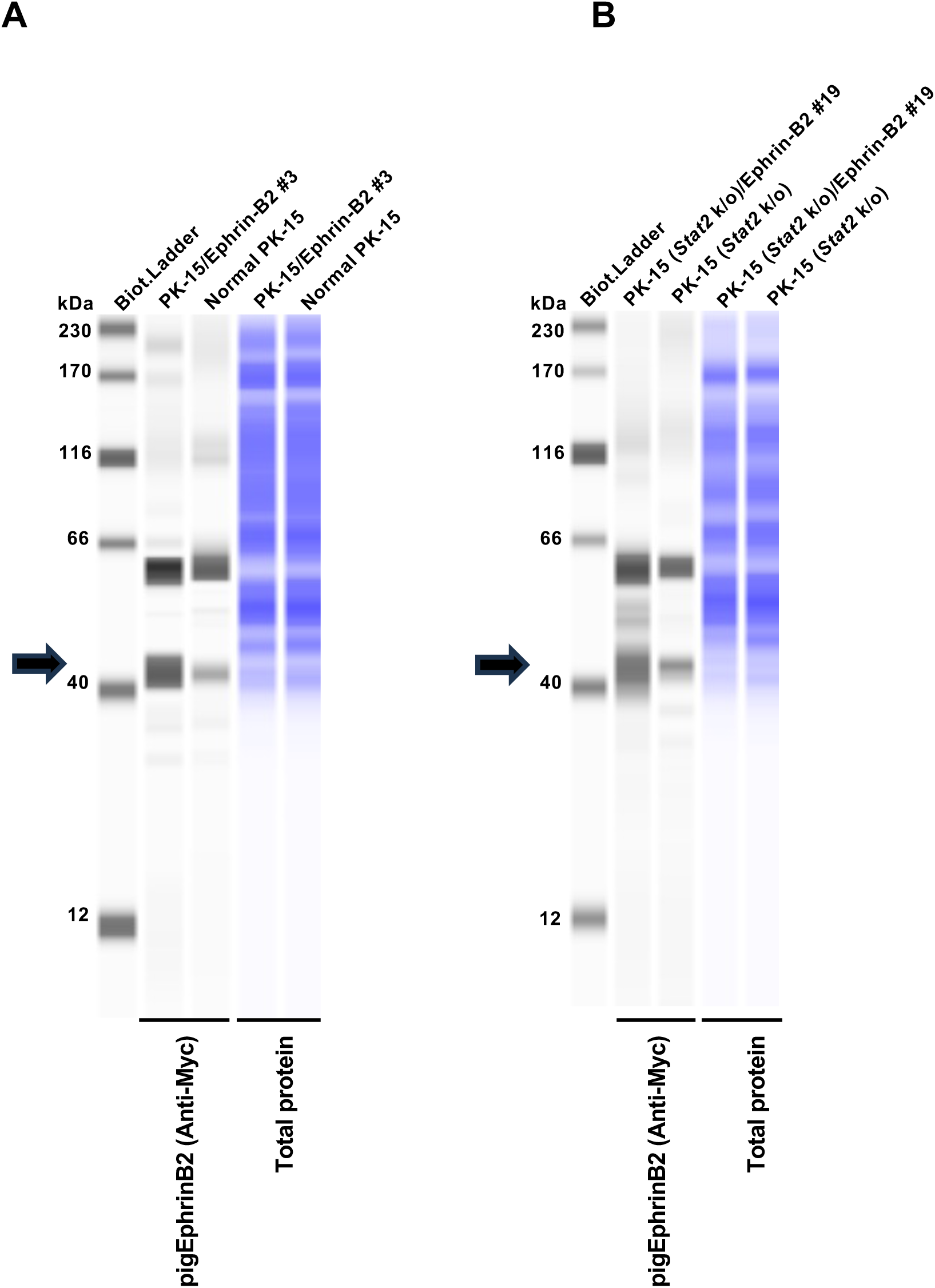
Generation of PK-15 cells stably expressing pig ephrin-B2. (a) Expression of Myc-tagged ephrin-B2 in PK-15 cells was evaluated by western blotting. The cellular lysate of normal PK-15 cells was used as a negative control (empty). Cellular lysates were probed with an anti-Myc antibody (left) and then re-probed with the Total Protein Detection Module (right). The black arrows indicate the size of Myc-tagged ephrin-B2. (b) Expression of Myc-tagged ephrin-B2 in PK-15 [*Stat2* knockout (*Stat2* k/o)] cells was evaluated by western blotting. The cellular lysate of unmodified PK-15 (*Stat2* k/o) cells was used as a negative control (empty). Cellular lysates were probed with an anti-Myc antibody (left) and then re-probed with the Total Protein Detection Module (right). The black arrows indicate the size of Myc-tagged ephrin-B2.

### 3.2 Ephrin-B2 Expression in PK-15 Cells Enhances NiV Pseudovirus Infectivity

We hypothesized that pig ephrin-B2 expression in PK-15 cells could enhance susceptibility to NiV infection. To address this, we generated HIV-1–and MLV-based vectors pseudotyped with NiV G and F proteins (termed NiV pseudovirus). Western blotting demonstrated efficient expression of NiV G (**Figure 3a**) and F proteins (**Figure 3b**) in the transfected cells.

**Figure 3.**
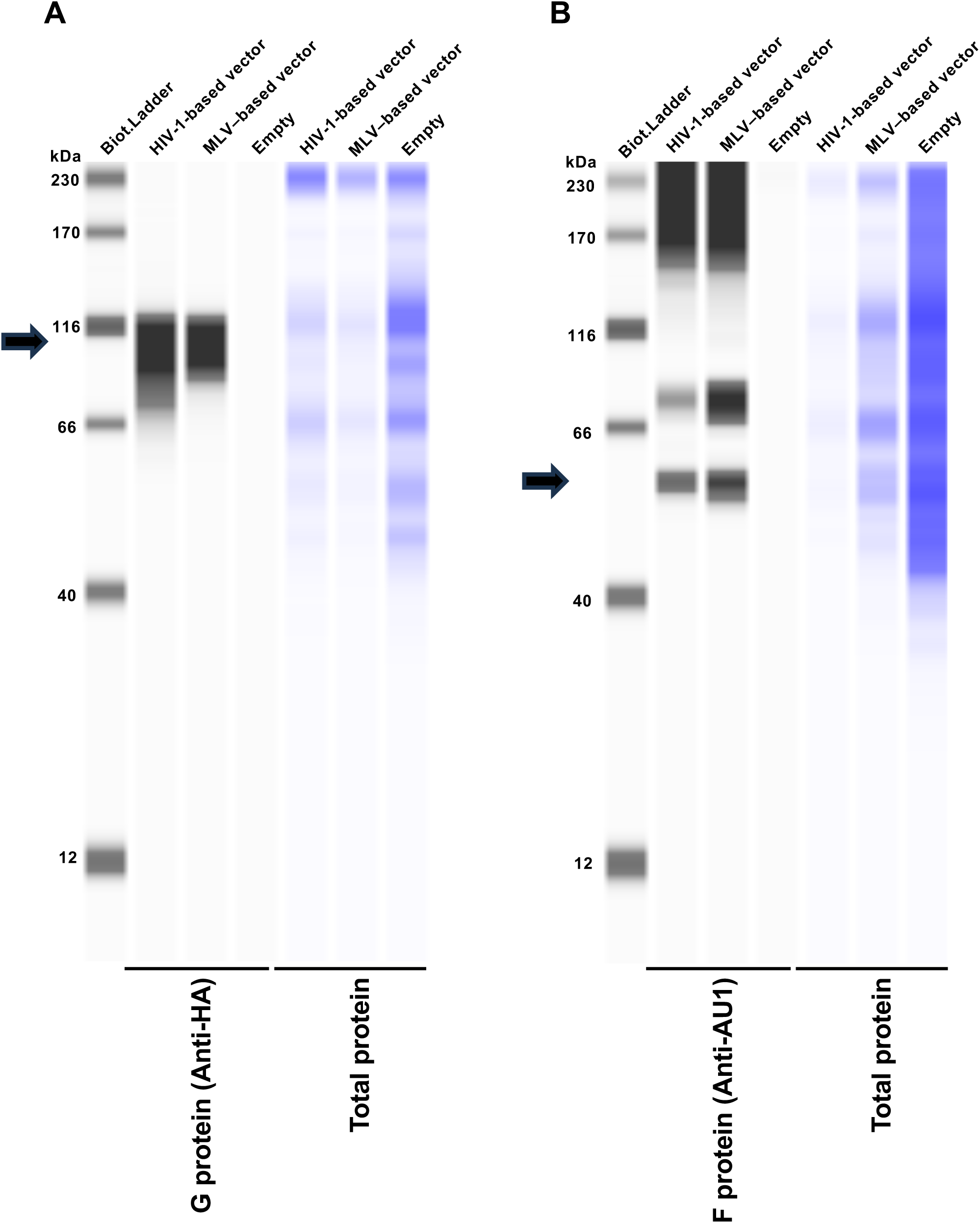
Production of Nipah virus (NiV) pseudovirus. (a) Expression of HA-tagged NiV G in Lenti-X 293T cells was evaluated by western blotting. The cellular lysate of Lenti-X 293T cells transfected with the pCAGGS empty plasmid was used as a negative control (empty). The black arrows indicate the sizes of the NiV G protein. (b) Expression of AU1-tagged NiV F in Lenti-X 293T cells was evaluated by western blotting. The cellular lysate of Lenti-X 293T cells transfected with the pCAGGS empty plasmid was used as a negative control (empty). The black arrows indicate the sizes of NiV F protein.

Next, we infected normal PK-15 and pig ephrin-B2–expressing PK-15 cells with the NiV pseudovirus. Infectivity was significantly enhanced by more than 1000-fold in PK-15/Ephrin-B2 #3 cells compared with that in normal PK-15 cells (**Figure 4a**).

**Figure 4.**
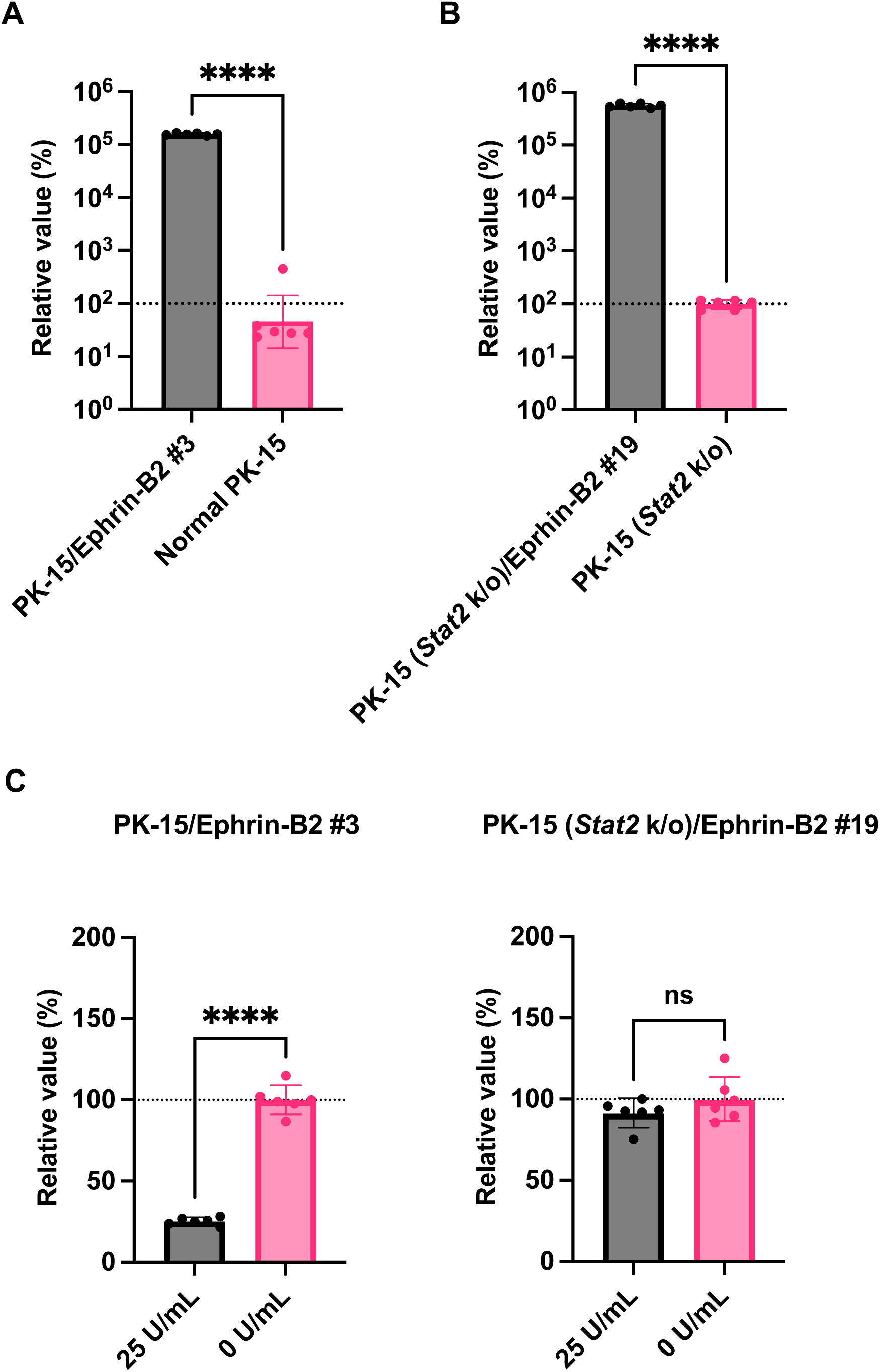
Stable ephrin-B2 expression in PK-15 cells enhances Nipah virus (NiV) pseudovirus infectivity. (a) PK-15 cells stably expressing ephrin-B2 (PK-15/Ephrin-B2 #3 cells) were infected with NiV pseudovirus. The luminescent signal was measured 2 days after infection. The relative value was calculated according to that in normal PK-15 cells. The results, presented as the mean and standard deviation of sextuplicate measurements from one assay, are representative of at least three independent experiments. Differences between PK-15/Ephrin-B2 #3 and normal PK-15 cells infected with NiV pseudovirus were examined using a two-tailed, unpaired Student’s *t*-test. **** *p* < 0.0001. (b) PK-15 [*Stat2* knockout (*Stat2* k/o)]/Ephrin-B2 #19 cells were infected with NiV pseudovirus. The luminescent signal was measured 2 days after infection. The relative value was calculated according to that in PK-15 (*Stat2* k/o) cells. The results, presented as the mean and standard deviation of sextuplicate measurements from one assay, are representative of at least three independent experiments. Differences between PK-15 (*Stat2* k/o)/Ephrin-B2 #19 and PK-15 (*Stat2* k/o) cells infected with NiV pseudovirus were examined using a two-tailed, unpaired Student’s *t*-test. **** *p* < 0.0001. (c) PK-15/Ephrin-B2 #3 and PK-15 (*Stat2* k/o)/Ephrin-B2 #19 cells were treated with 25 units/mL Universal Type I IFN or left untreated for 24 h and then infected with NiV pseudovirus. The luminescent signal was measured 2 days after infection. The results, presented as the mean and standard deviation of sextuplicate measurements from one assay, are representative of at least three independent experiments. The relative value was calculated according to that in PK-15 cells without Universal Type I IFN treatment. **** *p* < 0.0001, ns (not significant).

Type I IFN exerts antiviral effects by binding to IFNAR1 on the cellular surface, and STAT2 plays a critical role in this response. To avoid the inhibition of viral isolation by IFN, we recently generated a PK-15 cell line lacking *Ifnar1* or *Stat2* [22]. In this study, we tested the usefulness of PK-15 (*Stat2* k/o)/Ephrin-B2 #19 cells for investigating NiV pseudovirus infectivity in the presence of IFNs. Virus infectivity was higher in PK-15 (*Stat2* k/o)/Ephrin-B2 #19 cells lacking type I IFN than in PK-15 (*Stat2* k/o) cells (**Figure 4b**). We also observed a significant difference in infectivity in PK-15/Ephrin-B2 #3 with or without type I IFN treatment (**Figure 4c**). Conversely, virus infectivity in PK-15 (*Stat2* k/o)/Ephrin-B2 #19 cells was not affected by the addition of type I IFN (**Figure 4c**). These results suggest that Ephrin-B2 expression in PK-15 cells enhances NiV pseudovirus infectivity. Furthermore, in the presence of IFN or IFN-inducible substances in clinical samples, PK-15 (*Stat2* k/o) cells with stable ephrin-B2 expression could be more useful for efficient virus isolation than PK-15/Ephrin-B2 #3 cells.

### 3.3 No Enhanced Replication of Ephrin-B2–independent Viruses in PK-15/Ephrin-B2 #3 Cells

We then investigated the specificity of the enhancement in NiV pseudovirus infectivity in PK-15/Ephrin-B2 #3 cells. Specifically, we tested whether pig ephrin-B2 enhances the infection of an HIV-1–based lentiviral vector pseudotyped with VSV-G, the infection of which is mediated by endocytosis [24]. No difference in infection efficiency was observed between normal PK-15 and PK-15/Ephrin-B2 #3 cells (**Figure 5a**), suggesting that pig ephrin-B2 expression does not affect the infectivity of ephrin-B2–independent viruses. Therefore, stable expression of pig ephrin-B2 in PK-15 cells specifically promoted NiV pseudovirus infectivity.

**Figure 5.**
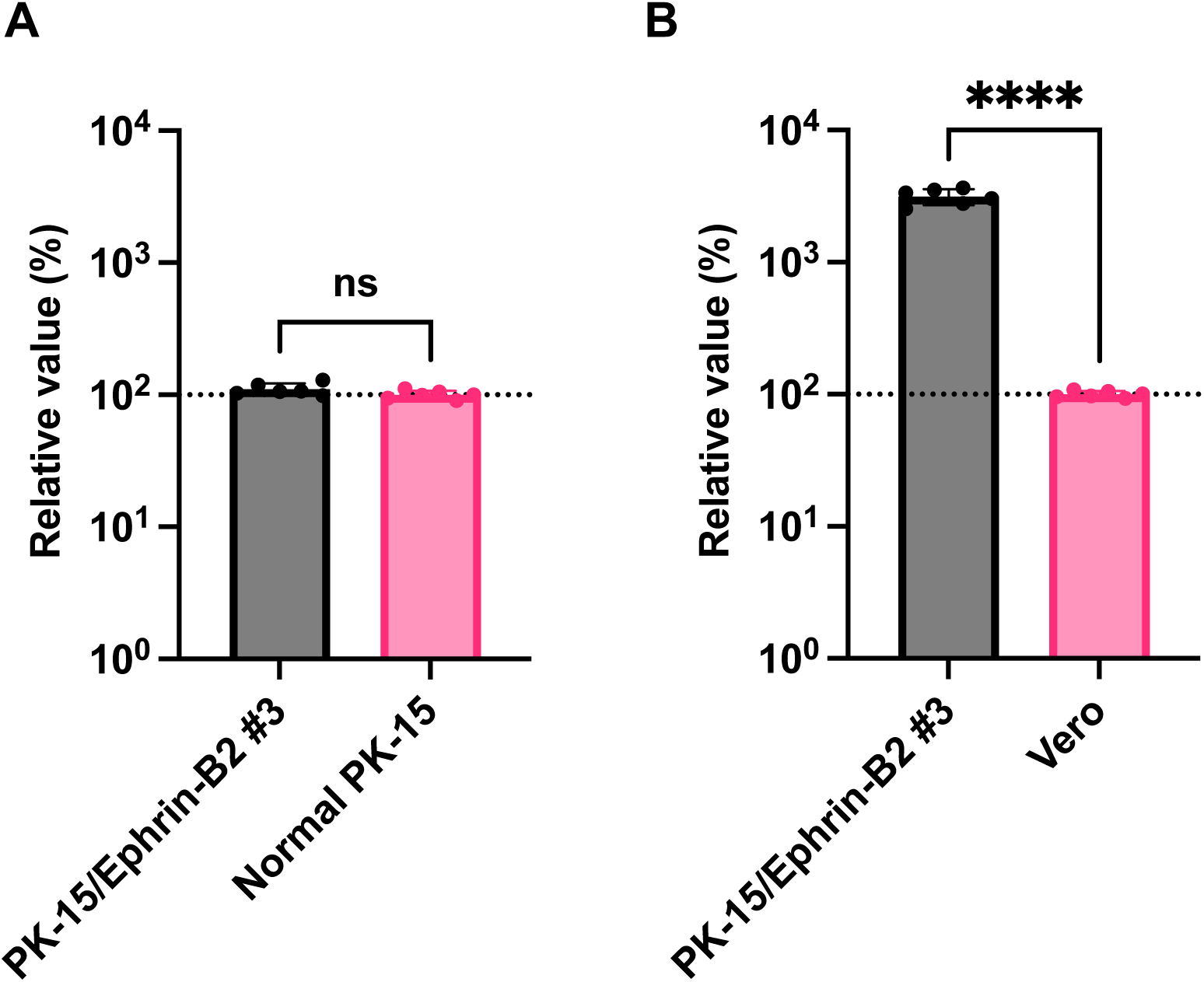
Nipah virus (NiV) pseudovirus more efficiently infected PK-15 cells stably expressing ephrin-B2 (PK-15/Ephrin-B2 #3 cells) than in Vero cells. (a) PK-15/Ephrin-B2 #3 and normal PK-15 cells were infected with HIV-1 vector pseudotyped with VSV-G, and the luminescent signal was measured 2 days after infection. The results, presented as the mean and standard deviation of sextuplicate measurements from one assay, are representative of at least three independent experiments. Differences between PK-15/Ephrin-B2 #3 and normal PK-15 cells were examined using a two-tailed, unpaired Student’s *t*-test. ns (not significant). (b) PK-15/Ephrin-B2 #3 or Vero E6 cells were infected with NiV pseudovirus, and the luminescent signal was measured 2 days after infection. The relative values were calculated in comparison to those in Vero E6 cells. The results, presented as the mean and standard deviation of sextuplicate measurements from one assay, are representative of at least three independent experiments. Differences between PK-15/Ephrin-B2 #3 and Vero E6 cells were examined using a two-tailed, unpaired Student’s *t*-test. **** *p* < 0.0001.

### 3.4 NiV Pseudovirus More Efficiently Infected PK-15/Ephrin-B2 #3 Cells than Vero Cells

Having demonstrated that NiV pseudovirus robustly infected PK-15/Ephrin-B2 #3 cells (**Figure 4a**), we hypothesized that PK-15/Ephrin-B2 #3 cells would be superior to other cell lines for NiV propagation. To investigate this, we compared NiV pseudovirus infectivity between PK-15/Ephrin-B2 #3 and Vero E6 cells, which WOAH recommends for NiV isolation. As TRIM5 in Vero cells blocks infection by HIV-1– based lentiviral vectors [25], we used an MLV-based retroviral vector pseudotyped with NiV G and F proteins. The result demonstrated that NiV pseudovirus infectivity was higher (>30-fold) in PK-15/Ephrin-B2 #3 cells than in Vero E6 cells (**Figure 5b**). Thus, PK-15/Ephrin-B2 #3 cells could be promising for NiV propagation.

### 3.5 PK-15/Ephrin-B2 #3 Cells Can Be Used for Neutralization Testing

We tested whether neutralization antibodies against NiV could block NiV pseudovirus infection in PK-15/Ephrin-B2 #3 cells. Neutralization tests illustrated that different dilutions of anti-NiV hyperimmune mouse ascitic fluid neutralized NiV pseudovirus in PK-15/Ephrin-B2 #3 cells (**Figure 6a**). The calculated NT_50_ of anti-NiV hyperimmune mouse ascitic fluid was 349.5-fold dilution. Conversely, ascitic fluid did not exert any effect against the HIV-1–based lentiviral vector pseudotyped with VSV-G in PK-15/Ephrin-B2 #3 cells (**Figure 6b**), highlighting the specificity of our assay.

**Figure 6.**
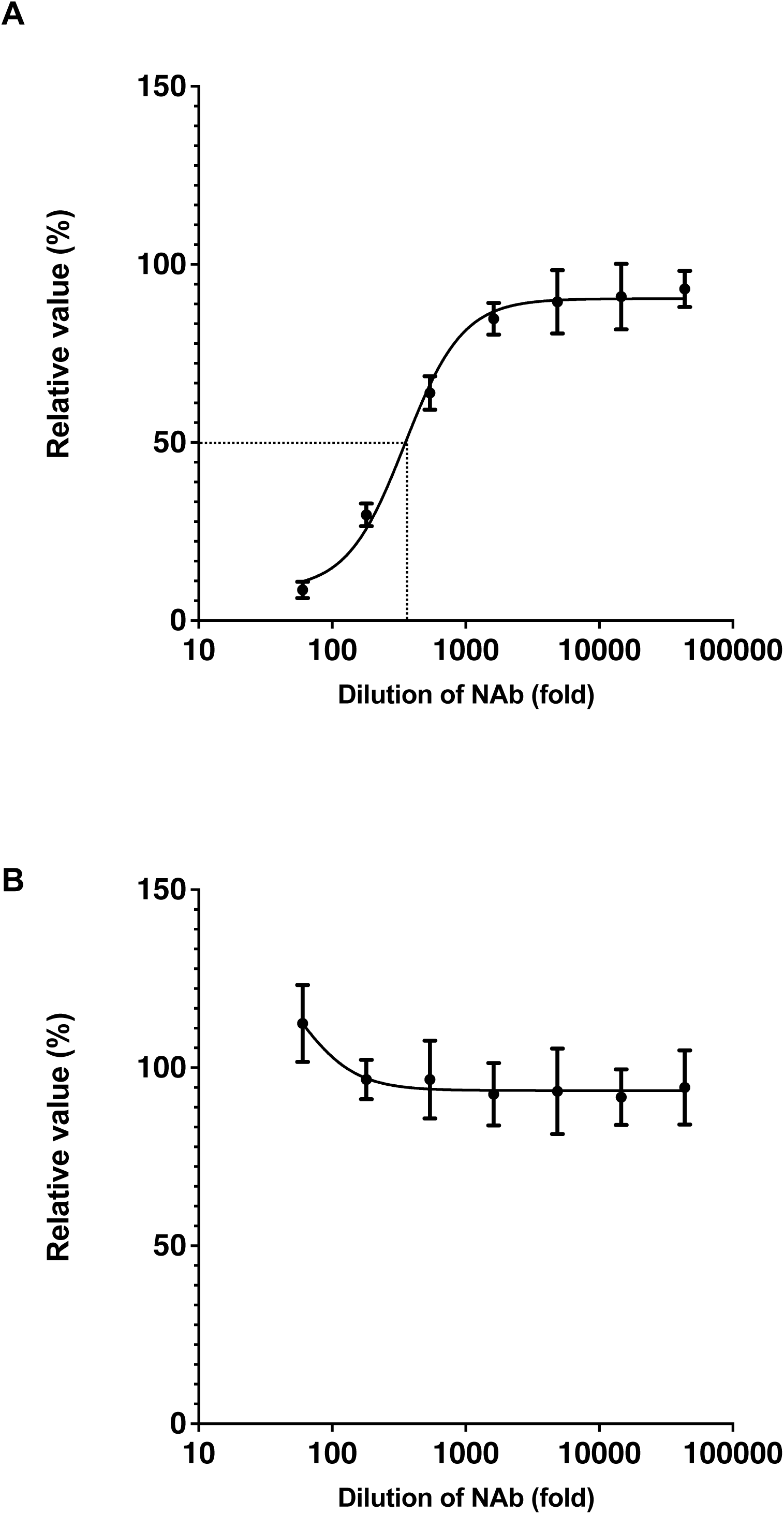
PK-15 cells stably expressing ephrin-B2 (PK-15/Ephrin-B2 #3 cells) can be used in neutralization assays. (a) NiV pseudovirus was mixed with serially diluted anti-Nipah virus (NiV) hyperimmune ascitic fluid for 1 h. The mixture was then added to PK-15/Ephrin-B2 #3 cells. The NT_50_ of anti-NiV hyperimmune ascitic fluid against NiV pseudovirus was measured 2 days after infection. The results, presented as the mean and standard deviation of sextuplicate measurements from one assay, are representative of at least three independent experiments. (b) HIV-1 vector pseudotyped with VSV-G was mixed with serially diluted anti-NiV hyperimmune ascitic fluid for 1 h. The mixture was then added to PK-15/Ephrin-B2 #3 cells. The NT_50_ of anti-NiV hyperimmune ascitic fluid against HIV-1 vector pseudotyped with VSV-G was measured 2 days after infection. The results, presented as the mean and standard deviation of sextuplicate measurements from one assay, are representative of at least three independent experiments.

## 4. Discussion

To date, no vaccines or drugs are approved for NiV infection in pigs or humans. Virus isolation from clinical specimens is essential for understanding the growth characteristics and pathogenicity of virus strains, and isolated viruses can be used to prepare vaccine antigens. In this study, we generated a porcine PK-15 cell line with stable pig ephrin-B2 expression to enhance NiV replication. The infectivity of NiV pseudovirus was significantly enhanced by >1000-fold in PK-15/Ephrin-B2 #3 cells compared with that in wild-type PK-15 cells, and NiV pseudovirus was markedly and specifically inhibited by an anti-NiV neutralizing antibody. We believe that PK-15/Ephrin-B2 #3 cells can improve the viral isolation efficiency of clinical samples from the respiratory or neurological organs of pigs infected with NiV. However, as our laboratory facilities are not capable of conducting biosafety level 4 (BSL-4) experiments, we used a pseudotyped virus instead of replication-competent NiV. Further studies are required to evaluate NiV replication efficiency in a BSL-4 laboratory.

Ephrin-B2 is a widely used receptor in henipaviruses, including NiV, Hendra virus, and Cedar virus [26]. As viruses exhibit optimal replication in their natural hosts, we believe that PK-15/Ephrin-B2 #3 cells can be an essential tool for isolating henipaviruses from pigs. Furthermore, we successfully generated PK-15 (*Stat2* k/o)/Ephrin-B2 #19 cells to isolate viruses in the presence of IFNs. To prove the usefulness of these cells, we will test the isolation efficiency of a variety of pig-derived viruses in future studies.

The lack of enhanced infectivity of lentiviral vectors pseudotyped with VSV-G in PK-15/Ephrin-B2 #3 cells confirmed that the observed enhanced infectivity of NiV pseudovirus in this cell line was strictly dependent on ephrin-B2–mediated entry. This specificity is crucial for diagnostic applications. Furthermore, we observed >30-fold higher NiV pseudovirus infectivity in PK-15/Ephrin-B2 #3 cells than in Vero cells, underscoring the higher sensitivity of the latter cell line for NiV infection. Additionally, some monoclonal antibodies, such as m102.4 and Hu1F5, have been tested in non-human primates, and they exhibited promising postexposure prophylactic and therapeutic effects [27, 28]. The neutralization of NiV pseudovirus in PK-15/Ephrin-B2 #3 cells by anti-NiV hyperimmune ascitic fluid validated the utility of the developed cells for neutralization assays in vaccine and therapeutic antibodies development.

Our study had some limitations. First, as previously discussed, confirmatory tests using replication-competent NiV to evaluate the replication efficiency of the virus in PK-15/Ephrin-B2 #3 cells in a BSL-4 laboratory are required. Second, the adverse effects of stable ephrin-B2 expression on the isolation or propagation of ephrin-B2–independent viruses should be considered. In this regard, we did not observe any adverse effect of stable ephrin-B2 expression on the infectivity of a lentiviral vector pseudotyped with VSV-G. Nevertheless, we must assess this possibility further in future studies.

In conclusion, we established a pig-derived PK-15 cell line stably expressing pig ephrin-B2, and NiV pseudovirus infectivity was significantly enhanced in these cells, suggesting a promising approach for isolating and propagating henipaviruses. We believe our cells will make significant contributions to both NiV research and preparedness for *Henipavirus* disease outbreaks in the pig population and assessments of *Henipavirus* transmission risk from pigs to humans.

## Acknowledgments

pMD2.G was a gift from Dr. Didier Trono. psPAX2-IN/HiBiT and pWPI-Luc2 were kind gifts from Dr. Kenzo Tokunaga. The authors thank Ms. Tomoko Nishiuchi, Ms. Miki Kawano, and the staff of CADIC, University of Miyazaki, for their assistance. This study was supported by the China Scholarship Council (to H.Z.). We additionally thank Enago (www.enago.com) for the English language review.

## Supporting information

**Supplementary Text 1. Synthesized DNA for generating a plasmid expressing pig ephrin-B2.** The coding sequence of pig ephrin-B2 was synthesized according to the amino acid sequence deposited in GenBank (accession number: ABV44489.1) with codon optimization to pig cells. The start and stop codons are underlined.

**Supplementary Text 2. Synthesized DNA for generating a plasmid expressing Nipah virus (NiV) G protein (A) and F protein (B).** The coding sequences of NiV G and F proteins were synthesized according to the amino acid sequences deposited in GenBank (accession numbers: AAK29088.1 and AAK29087.1). The start and stop codons are underlined.

## List of abbreviations

NiV: Nipah virus
PK-15/Ephrin-B2 cells: PK-15 cells stably expressing pig-derived ephrin-B2
IFN: interferon
IFNAR: interferon alpha and beta receptor subunit 1
*Stat2* k/o cells: *Stat2* knockout cells
PK-15 (*Stat2* k/o)/Ephrin-B2 #19 cells: *Stat2* knockout cells expressing pig-derived ephrin-B2

## References

1. McLean RK, Graham SP. The pig as an amplifying host for new and emerging zoonotic viruses. One Health 2022; 14: 100384.

2. Kulkarni DD, Tosh C, Venkatesh G, Senthil Kumar D. Nipah virus infection: current scenario. Indian J Virol 2013; 24(3): 398–408.

3. Chua KB, Bellini WJ, Rota PA, et al. Nipah virus: a recently emergent deadly paramyxovirus. Science 2000; 288(5470): 1432–5.

4. Parashar UD, Sunn LM, Ong F, et al. Case-control study of risk factors for human infection with a new zoonotic paramyxovirus, Nipah virus, during a 1998-1999 outbreak of severe encephalitis in Malaysia. J Infect Dis 2000; 181(5): 1755–9.

5. Chowdhury S, Khan SU, Crameri G, et al. Serological evidence of henipavirus exposure in cattle, goats and pigs in Bangladesh. PLoS Negl Trop Dis 2014; 8(11): e3302.

6. Bonaparte MI, Dimitrov AS, Bossart KN, et al. Ephrin-B2 ligand is a functional receptor for Hendra virus and Nipah virus. Proc Natl Acad Sci U S A 2005; 102(30): 10652–7.

7. Negrete OA, Levroney EL, Aguilar HC, et al. EphrinB2 is the entry receptor for Nipah virus, an emergent deadly paramyxovirus. Nature 2005; 436(7049): 401–5.

8. Negrete OA, Wolf MC, Aguilar HC, et al. Two key residues in ephrinB3 are critical for its use as an alternative receptor for Nipah virus. PLoS Pathog 2006; 2(2): e7.

9. Bossart KN, Tachedjian M, McEachern JA, et al. Functional studies of host-specific ephrin-B ligands as Henipavirus receptors. Virology 2008; 372(2): 357–71.

10. AbuBakar S, Chang LY, Ali AR, Sharifah SH, Yusoff K, Zamrod Z. Isolation and molecular identification of Nipah virus from pigs. Emerg Infect Dis 2004; 10(12): 2228–30.

11. Chua KB, Goh KJ, Wong KT, et al. Fatal encephalitis due to Nipah virus among pig-farmers in Malaysia. Lancet 1999; 354(9186): 1257–9.

12. Mohd Nor MN, Gan CH, Ong BL. Nipah virus infection of pigs in peninsular Malaysia. Rev Sci Tech 2000; 19(1): 160–5.

13. Wong KT, Shieh WJ, Kumar S, et al. Nipah virus infection: pathology and pathogenesis of an emerging paramyxoviral zoonosis. Am J Pathol 2002; 161(6): 2153–67.

14. DeBuysscher BL, Scott DP, Rosenke R, Wahl V, Feldmann H, Prescott J. Nipah Virus Efficiently Replicates in Human Smooth Muscle Cells without Cytopathic Effect. Cells 2021; 10(6).

15. Ang LT, Nguyen AT, Liu KJ, et al. Generating human artery and vein cells from pluripotent stem cells highlights the arterial tropism of Nipah and Hendra viruses. Cell 2022; 185(14): 2523–41 e30.

16. Elvert M, Sauerhering L, Maisner A. Cytokine Induction in Nipah Virus-Infected Primary Human and Porcine Bronchial Epithelial Cells. J Infect Dis 2020; 221(Suppl 4): S395–S400.

17. Elvert M, Sauerhering L, Heiner A, Maisner A. Isolation of Primary Porcine Bronchial Epithelial Cells for Nipah Virus Infections. Methods Mol Biol 2023; 2682: 103–20.

18. Muller M, Fischer K, Woehnke E, et al. Analysis of Nipah Virus Replication and Host Proteome Response Patterns in Differentiated Porcine Airway Epithelial Cells Cultured at the Air-Liquid Interface. Viruses 2023; 15(4).

19. Sauerhering L, Zickler M, Elvert M, et al. Species-specific and individual differences in Nipah virus replication in porcine and human airway epithelial cells. J Gen Virol 2016; 97(7): 1511–9.

20. World Organization of Animal Health. NIPAH AND HENDRA VIRUS DISEASES. 2022. Available from: https://www.woah.org/fileadmin/Home/eng/Health_standards/tahm/3.01.15_NIPAH_HENDRA.pdf

21. Sadler AJ, Williams BR. Interferon-inducible antiviral effectors. Nat Rev Immunol 2008; 8(7): 559–68.

22. Shofa M, Saito A. Generation of porcine PK-15 cells lacking the Ifnar1 or Stat2 gene to optimize the efficiency of viral isolation. PLoS One 2023; 18(11): e0289863.

23. Bradel-Tretheway BG, Zamora JLR, Stone JA, Liu Q, Li J, Aguilar HC. Nipah and Hendra Virus Glycoproteins Induce Comparable Homologous but Distinct Heterologous Fusion Phenotypes. J Virol 2019; 93(13).

24. Albertini AA, Baquero E, Ferlin A, Gaudin Y. Molecular and cellular aspects of rhabdovirus entry. Viruses 2012; 4(1): 117–39.

25. Berthoux L, Sebastian S, Sokolskaja E, Luban J. Cyclophilin A is required for TRIM5alpha-mediated resistance to HIV-1 in Old World monkey cells. Proc Natl Acad Sci U S A 2005; 102(41): 14849–53.

26. Laing ED, Navaratnarajah CK, Cheliout Da Silva S, et al. Structural and functional analyses reveal promiscuous and species specific use of ephrin receptors by Cedar virus. Proc Natl Acad Sci U S A 2019; 116(41): 20707–15.

27. Playford EG, Munro T, Mahler SM, et al. Safety, tolerability, pharmacokinetics, and immunogenicity of a human monoclonal antibody targeting the G glycoprotein of henipaviruses in healthy adults: a first-in-human, randomised, controlled, phase 1 study. Lancet Infect Dis 2020; 20(4): 445–54.

28. Zeitlin L, Cross RW, Woolsey C, et al. Therapeutic administration of a cross-reactive mAb targeting the fusion glycoprotein of Nipah virus protects nonhuman primates. Sci Transl Med 2024; 16(741): eadl2055.

