## Supplementary Text for "Development of a Porcine Cell Line Stably Expressing Ephrin-B2 for Nipah Virus Research and Diagnostic Testing"

**Supplementary Text 1. Synthesized DNA for generating a plasmid expressing pig EphrinB2**

ATGGCAGCTAGAAGGGACAGCGTCTGGAAGTATTGTTGGGGTGTTTTGATGGTGCTCTGT
CGCACAGCAATTTCAAGAAGTATAGTTCTTGAACCGATCTATTGGAATAGCTCTAACTCAA
AGTTCCTTCCTGGACAGGGACTGGTCCTTTACCCACAGATCGGGGATAAGCTGGATATAA
TTTGCCCGAAGGTCGACAGTAAGACCGTGGGACAATATGAGTATTACAAAGTGTATATGG
TGGATAAGGATCAGGCCGATCGCTGCACGATTAAGAAAGAGAACACGCCTCTCCTGAATT
GTGCTAGACCTGACCAGGATGTCAAATTCACAATCAAGTTTCAAGAATTTAGCCCGAACC
TGTGGGGGCTCGAATTTCAGAAGAACAGGGATTACTATATCATTTCAACGAGCAACGGTT
CCCTCGAGGGATTGGACAATCAAGAAGGGGGCGTCTGCCAAACACGCGCGATGAAAATA
CTGATGAAAGTGGGACAAGACGCCAGCAGTGCCGGTTCTACCAGACATAACGAACCGAC
AAGACGGCCTGAACTCGAGGCGGGCACGAACGGTCGCTCTAGTACGACGAGTCCTTTTG
TCAAGCCAAATCCTGGGTCCAGTACAGATGGGAACAGCGCGGGCCATAGTGGTAATAATA
TTCTGGGTTCCGAAGTTGCGCTCTTTGCCGGCATCGCGTCAGGCTGCATCATTTTCATTGT
GATTATTATCACACTCGTTGTCCTCTTGCTGAAATATAGGAGGAGGCACCGGAAGCATTCC
CCCCAACACACTGCCACCCTCTCTATCTCAACGTTGGCTACTCCGAAGAGGGGTGGGAAT
AACAATGGCAGCGAGCCTTCCGACATTATTATTCCATTGAGAACGGCAGATAGTGTGTTCT
GCCCGCACTATGAGAAGGTCAGCGGTGACTATGGACATCCAGTTTACATAGTGCAAGAAA
TGCCCCCGCAGTCCCCTGCCAATATTTACTATAAAGTCGAGCAGAAGCTTATTTCCGAAGA
AGATCTCTAA

**Supplementary Text 2. Synthesized DNA for generating a plasmid expressing Nipah virus (NiV) G protein (A) and F protein (B).**

### A. Nipah virus G protein

<u>ATG</u>CCAGCGGAGAATAAGAAAGTGAGATTCGAGAATACTACGAGCGACAAAGGCAAGAT
ACCCAGCAAGGTAATAAAAAGCTACTATGGTACAATGGACATAAAAAAAATAAACGAAG
GGCTCCTGGACAGCAAGATCCTTTCAGCTTTTAACACGGTGATAGCACTCTTGGGCTCAA
TAGTGATTATCGTGATGAACATAATGATTATTCAGAATTACACTCGGAGTACTGATAATCAA
GCAGTCATAAAGGATGCTCTTCAGGGCATCCAACAACAGATAAAAGGCCTTGCCGACAA
AATCGGTACAGAAATTGGACCAAAAGTCTCACTTATAGACACCTCCTCAACAATAACCAT
CCCTGCTAATATTGGGCTCCTCGGCAGTAAGATCTCTCAGTCAACAGCGAGCATCAATGA
AAATGTGAACGAAAAGTGCAAGTTCACTCTGCCACCCCTGAAGATTCATGAGTGTAACAT
AAGCTGTCCGAATCCACTCCCTTTTCGGGAGTATAGGCCGCAAACTGAGGGAGTGAGCA
ACCTGGTGGGATTGCCGAACAACATCTGCTTGCAAAAGACGAGTAATCAGATTCTGAAG
CCGAAACTCATATCCTACACACTGCCAGTTGTCGGACAGAGTGGTACGTGTATCACCGAT
CCTCTTCTGGCGATGGATGAAGGGTATTTTGCATATTCCCATCTGGAACGCATCGGCTCCT
GCAGTCGAGGGGTATCCAAGCAGCGAATCATTGGTGTCGGCGAAGTGTTGGATCGAGGA
GATGAAGTGCCAAGTTTGTTCATGACGAACGTGTGGACCCCGCCAAATCCCAACACCGT
TTACCACTGTTCAGCTGTTTACAATAACGAGTTCTACTACGTTCTCTGTGCAGTGTCAACC
GTGGGCGACCCAATCCTTAATTCTACTTACTGGAGCGGGTCCCTTATGATGACAAGGCTC
GCCGTGAAACCGAAGAGTAACGGAGGGGGTTATAATCAACATCAGCTCGCCCTTCGCTCT
ATAGAAAAGGGCAGATACGATAAGGTTATGCCTTACGGTCCTAGCGGTATCAAGCAAGGA
GATACCCTTTATTTCCCCGCTGTAGGTTTTCTCGTCCGCACGGAGTTCAAATATAATGATTC
CAACTGTCCAATTACTAAGTGCCAATACTCCAAACCCGAGAATTGTAGATTGAGTATGGG
GATACGGCCCAACAGCCATTACATCCTGAGATCTGGGTTGCTGAAATACAACCTGTCAGA
TGGCGAGAACCCTAAAGTGGTCTTTATTGAGATCTCAGATCAGCGGTTGTCAATCGGGAG
TCCCAGCAAGATCTATGATTCTCTCGGACAGCCGGTCTTCTACCAGGCAAGCTTTTCATGG
GACACCATGATCAAATTTGGTGACGTTCTTACGGTAAATCCGCTTGTGGTAAACTGGCGC
AATAATACTGTGATCAGTCGGCCAGGTCAATCTCAATGCCCGCGCTTCAATACATGCCCCG
AGATCTGTTGGGAAGGTGTATACAACGACGCATTCCTGATCGACAGGATTAACTGGATAA
GCGCCGGAGTTTTCCTTGACTCAAACCAAACTGCTGAAAACCCAGTTTTCACAGTTTTCA
AAGACAACGAAATCTTGTATCGGGCACAGTTGGCCTCAGAGGATACGAATGCTCAAAAA
ACAATCACAAACTGTTTCCTCCTTAAGAACAAGATCTGGTGCATCAGCTTGGTAGAGATA
TACGACACCGGTGACAACGTTATCCGGCCAAAACTTTTCGCCGTAAAAATACCCGAACAA
TGCACGTATCCCTATGATGTCCCCGACTATGCT<u>TAG</u>

**B. Nipah virus F protein**

<u>ATG</u>GTAGTGATTCTTGACAAAAGGTGTTACTGCAATCTTCTTATTCTTATACTTATGATCTC
CGAATGCTCTGTGGGAATCCTGCACTACGAAAAGCTCAGTAAGATTGGGCTCGTCAAGG
GAGTTACCAGGAAGTACAAGATCAAATCTAATCCACTCACAAAGGACATAGTTATTAAGA
TGATTCCCAACGTCTCCAATATGTCACAGTGCACAGGGAGTGTAATGGAAAATTACAAAA
CAAGACTCAATGGCATCTTGACGCCGATAAAGGGTGCGCTTGAGATATATAAAAATAACA
CTCACGACTTGGACTACAAGGATGATGACGATAAGGTTGGAGACGTGAGATTGGCCGGA
GTTATTATGGCCGGTGTCGCTATTGGTATCGCGACCGCGGCCCAGATTACGGCAGGAGTAG
CGCTGTACGAGGCCATGAAGAATGCGGACAATATCAACAAACTCAAGAGTTCCATAGAA
AGTACTAATGAGGCTGTTGTGAAACTGCAAGAGACAGCTGAGAAGACTGTCTATGTATTG
ACCGCGTTGCAAGACTATATTAACACTAACCTCGTTCCGACAATAGACAAGATAAGTTGTA
AGCAAACTGAGCTTAGCTTGGACCTGGCATTGTCCAAGTACCTTAGCGATCTGCTGTTTG
TATTTGGTCCGAACTTGCAGGACCCAGTATCTAACTCAATGACTATTCAGGCTATATCCCA
GGCTTTCGGCGGGAACTACGAAACACTTTTGAGGACGCTCGGGTACGCAACTGAGGACT
TTGATGATCTCCTGGAGTCTGATTCTATTACGGGACAGATAATCTACGTTGATCTCAGTTCT
TATTATATAATCGTGCGCGTCTATTTCCCGATTCTCACTGAAATTCAACAAGCGTACATACA
GGAGCTGCTGCCCGTGAGCTTCAACAATGATAACAGCGAATGGATCAGTATCGTGCCAAA
CTTCATCTTGGTCAGAAACACCTTGATTAGTAACATAGAGATCGGATTTTGTCTGATCACT
AAGCGCAGTGTCATCTGCAATCAAGATTACGCAACTCCAATGACCAACAATATGAGAGAA
TGTTTGACTGGGTCCACAGAAAAATGTCCAAGAGAGCTTGTGGTGTCTTCCCACGTGCC
GCGGTTTGCCTTGAGCAACGGAGTACTTTTCGCGAATTGTATTTCAGTCACCTGCCAATGT
CAGACAACGGGTCGAGCAATATCACAGTCAGGGGAGCAAACCCTGCTGATGATAGATAAT
ACCACCTGTCCTACTGCCGTGCTCGGCAATGTCATCATTTCTTTGGGCAAATACCTTGGAT
CAGTTAATTACAATTCAGAGGGTATTGCAATAGGCCCTCCGGTCTTCACGGATAAGGTTGA
TATTTCTTCACAGATTAGTAGTATGAATCAGTCTCTGCAGCAGTCTAAGGACTATATAAAA
GAGGCACAACGCCTCCTTGACACCGTGAACCCTTCACTGATAAGCATGCTTAGCATGATA
ATTCTGTATGTTTTGAGTATCGCAAGTCTTTGTATAGGACTTATTACGTTTATAAGTTTTATT
ATAGTTGAGAAAAAAAGAAATACCTACTCTAGATTGGAGGACAGGAGAGTGAGACCTAC
TTCATCCGGAGATCTCTACTACATTGGCACCGACACCTACAGGTATATT<u>TGA</u>
